# ZEB2 regulates the development of CD11c+ atypical B cells

**DOI:** 10.1101/2022.09.01.506173

**Authors:** Xin Gao, Qian Shen, Jonathan A. Roco, Katie Frith, C. Mee Ling Munier, Maxim Nekrasov, Becan Dalton, Jin-Shu He, Rebecca Jaeger, Matthew C. Cook, John J. Zaunders, Ian A. Cockburn

## Abstract

CD11c^+^ atypical B cells (ABC) are an alternative memory B cell lineage identified both in normal immune responses as well as pathogenic responses in autoimmunity. While it is clear that ABCs have a distinct transcriptional program, the factors that direct this program have not been identified. Here, we generated a human tonsil single-cell RNA-seq dataset and identified candidate transcription factors associated with the ABC population. We selected 8 of these transcription factors for further analysis based on their conserved expression in mouse ABC bulk RNA-seq datasets. Using an optimized CRSPR-Cas9 knockdown method we found that only zinc finger E-box binding homeobox 2 (*Zeb2*) knock-out impaired ABC formation. To assess the role of Zeb2 in ABC formation in vivo we used Zeb2^fl/fl^ mice crossed to a CD23^Cre^ line. Germinal center and plasma cell responses in these mice after *Plasmodium* sporozoite immunization were largely unaltered but we observed a specific defect in ABC formation. We further determined that *ZEB2* haploinsufficient Mowat Wilson syndrome patients also have decreased circulating ABCs in the blood, supporting a role for this transcription factor in humans as well as mice. In sum, we identified Zeb2 as a key TF governing the formation of ABCs.

## Introduction

Following the resolution of infection, populations of memory lymphocytes, including B cells persist. Classical memory B cells (cMBCs) express high levels of CD27 and rapidly secrete antibody in response to restimulation. However non-classical memory B cells have also been identified in a variety of conditions. A population of FCRL4 expressing B cells was first identified in the tonsils of individuals undergoing tonsillectomy (1). Phenotypically similar FCRL4^+^ CD21^−^ CD27^−^ cells were subsequently found at elevated frequencies in the blood of HIV infected individuals (2). Similarly, individuals exposed to chronic malaria infection also had prominent populations of CD11c^+^ CD21^−^ CD27^−^ B cells (3–8). Collectively these cells associated with chronic infection were designated “atypical” B cells (ABCs). A related population of IgD^−^ CD27^−^ CXCR5^−^ CD21^−^ double negative 2 (DN2) B cells has also been identified in a variety of autoimmune conditions and their number generally correlates with disease severity (9–11). In animal models older mice were found to have a large population of CD11c^+^ B cells which were described as age-associated B cells (12, 13).

Recently it has become apparent that these heterogenous cells identified in a variety of autoimmune and infectious disease conditions nonetheless share a similar transcriptional signature (7). Moreover, while originally associated with chronic immune conditions or infection, ABCs have also been seen to respond to acute viral infections such as SARS-CoV-2 or primary vaccination with yellow fever, vaccinia virus vaccine or attenuated *Plasmodium* sporozoites (8, 14, 15). Extending this finding using single-cell RNA-seq (scRNA-seq) we found that ABCs were abundant not only in malaria exposed individuals, but also non-exposed European donors, but were undercounted due to the fact that many such cells retained expression of CD21 and CD27 (8).

While single cell transcriptomic data and associated trajectory analyses suggest that ABCs are part of an alternative B cell lineage, the key signals and transcription factors (TFs) that drive the formation of these cells are incompletely understood. IFNγ inducible T-box transcription factor 21 (Tbet) has been frequently found to be expressed on many alternative lineage cells in autoimmune and infectious disease conditions (5, 10, 16). Moreover ,the number of circulating CD11c^+^ CD21^low^ ABCs was low in one patient with inherited Tbet deficiency (17). However in animal models of systemic lupus erythematosus (SLE) Tbet has been found to be dispensable for disease progression and the formation of CD11c^+^ CD11b^+^ ABCs (18, 19). Moreover, in an *E. muris* model of infection CD19^hi^ CD11c^+^ ABCs formed normally in Tbet knockout animals (20).

Here we find, using a variety of human and mouse transcriptomic datasets, that alternative lineage B cells are consistently associated with the upregulation of several transcription factors including ZEB2. We developed a CRISPR/Cas9 knockdown screening protocol to ablate individual TFs and assess ABC formation in vitro and in vivo. In this screen Zeb2 was the only TF required for ABC development. We further confirmed this result using *Zeb2*^fl/fl^*Cd23*^Cre/+^ mice. Finally, to determine if there was a requirement for ZEB2 in ABC development in humans we examined the blood of *ZEB2* haplo-insufficient Mowat Wilson syndrome (MWS) patients who showed a specific defect in ABC formation.

## Results and Discussion

### Single cell RNA-seq of human tonsils identifies TFs associated with the ABC lineage

To extend our previous analysis of ABCs in the blood of healthy donors to lymphoid tissues we analysed B cells from the tonsils of two donors by single cell-RNA seq. To correlate transcriptomic data with surface marker expression we included a panel of 14 barcoded antibodies for cellular indexing of transcriptomes and epitopes by sequencing (CITE-seq) analysis. These antibodies (see Key Resource Table) were specific for surface markers that have previously been suggested to delineate different memory B cell subsets (21). Because tonsillar B cells are dominated by IgD^+^ naïve cells that were of limited interest to us, we sorted three samples from each donor (i) total tonsil B cells (ii) CD10^+^ IgD^−^ germinal center (GC) tonsillar B cells and (iii) CD10-IgD^−^ memory B cells (**Figure 1A; Figure S1A**). This allowed us to focus on populations of interest, without losing information about the relative frequencies of each population. In addition to our extended CITE-seq analysis we performed VDJ sequencing on the B cells to determine mutational frequencies.

**Figure 1.**
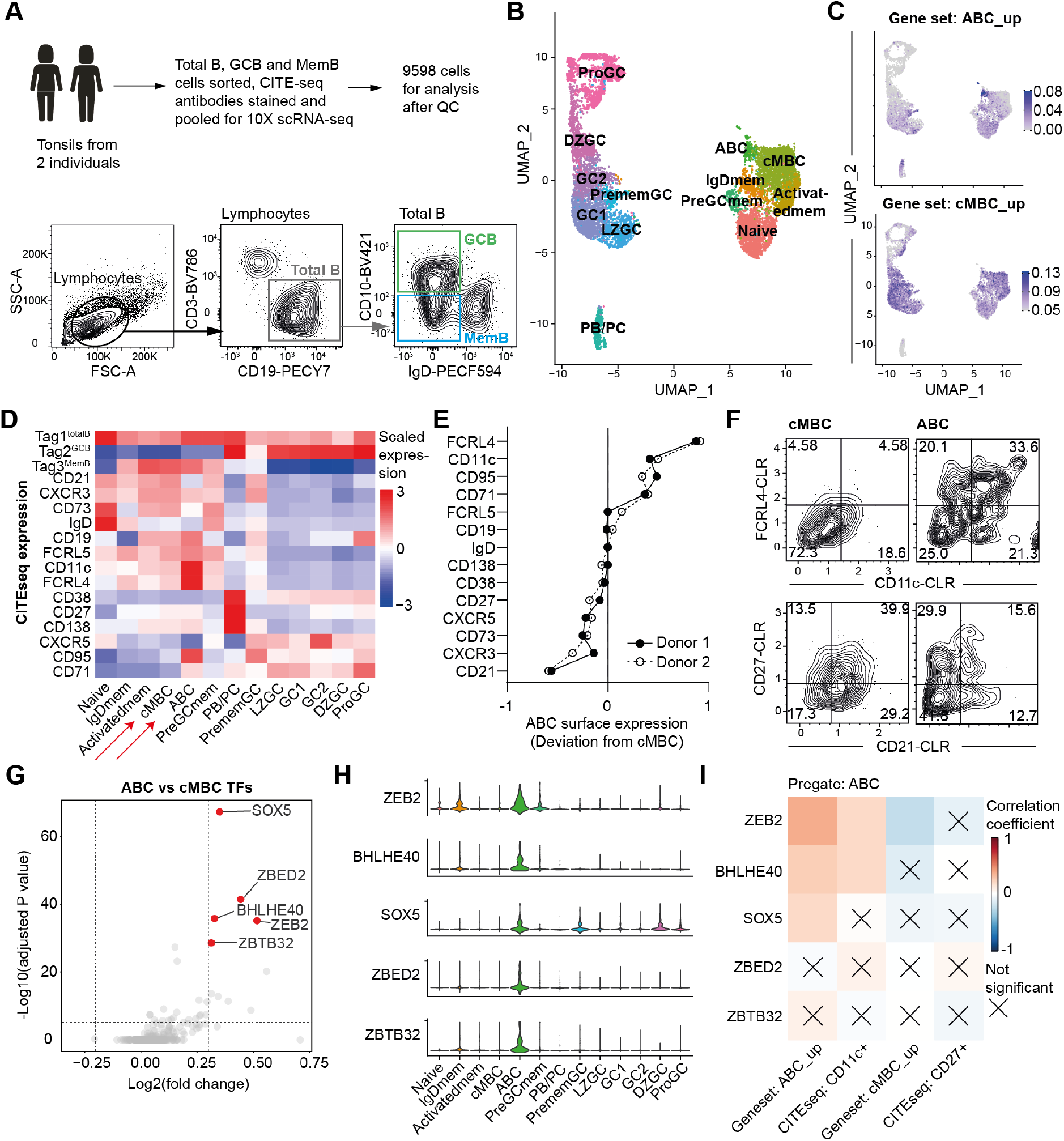
Single cell RNA-seq of human tonsils identifies TFs associated with the ABC lineage. Human tonsil total B cells, GC B cells and memory B cells were sorted, labelled by CITE-seq antibodies and mixed at 1:1:1 ratio for 10X scRNA-seq. The data was processed through Cellranger pipeline and analyzed by Seurat package. (A) Experiment design and gating strategy for sorting. (B) UMAP of B cells with identified clusters. (C) Feature plot showing the average expression of ABC and cMBC gene sets. (D) Heatmap of surface marker expression of all identified clusters by CITE-seq. (E) Statistics showing the relative expression of indicated surface markers in ABC comparing to cMBC. The deviation values were calculated as ABC CLR median value minus cMBC CLR median value. (F) Representative FACS plot showing the expression of indicated genes by CITE-seq. (G) Volcano plot comparing the expressions of TFs in ABC and cMBC, the top5 TFs were highlighted. (H) Stacked violin plot showing the expression of the top5 TFs in all clusters. (I) Pearson correlation tests between TFs expression and indicated features in each cell in ABC cluster were performed and the summarized as heatmap.

Consistent with previous scRNA-seq analyses of lymphoid tissues (22, 23), tonsillar B cells formed three distinct major clusters – a population of antibody secreting cells (plasmablast/plasmacell, PB/PC), a GC supercluster divided into light zone (LZ), dark zone (DZ), proliferating cells (ProGC) and unassigned GC1, GC2, and finally a supercluster of more quiescent cells that were a mix of naïve and memory cells (**Figure 1B; Figure S1A**). In addition to naïve cells and classical memory B cells (cMBC) this latter cluster included IgD^+^ memory cells, and some Pre-GC memory cells that are enriched with LZ genes such as CD83 (**Figure S1B**). A population of cells expressing *FOS, JUN* and *CD69* was observed, akin to the activated memory B cells in blood observed by us and others (**Figure S1B**) (8, 24). ABCs could be identified as a distinct peninsular within the memory/naïve cluster that was enriched for the expression of previously identified ABC genes and down regulation of cMBC genes (**Figure 1C**) (25). Cells in this ABC cluster also had the highest surface expression of FCRL4, CD11c and CD19 by CITE-seq analysis (**Figure 1D**). Data has been conflicting on whether ABC formation is dependent on the GC (20, 26), therefore we performed VDJ analysis on identified clusters and we found ABCs were not notably different from cMBCs with respect to their antibody isotype usage (**Figure S1C**). Moreover, they had only marginally lower frequencies of somatic hypermutation (SHM) compared to cMBCs and GC B cells (**Figure S1D**), though SHM inversely correlated with the strength of expression of ABC genes (**Figure S1E**). These results suggest the majority of tonsillar ABCs are GC-experienced.

Studies in ABC biology have used different surface markers (such as FCRL4, CD21, CD27 and CD11c) to identify ABCs whereas it remains unclear whether these markers have similar power to delineate ABCs from other B cell populations. Therefore, we examined all the surface markers in our CITE-seq dataset and found FCRL4, CD21 and CD11c were most useful to delineate ABC from cMBC in tonsils in both donors (**Figure 1E, Figure S1F**). Interestingly, we also found GC B cell markers CD95 and CD71 were upregulated in tonsillar ABCs. Closer examination of the surface phenotype of cells in the ABC cluster by the top markers FCRL4 vs CD11c and conventionally used CD27 vs CD21 revealed significant heterogeneity (**Figure 1F**). The ABC population was 33.6% FCRL4^+^ CD11c^+^ (compared to 4.58% of cMBCs) and 41.8% CD21^−^ CD27^−^ (compared to 17.3% of cMBCs). Collectively, these data suggest although FCRL4, CD21 and CD11c are the better surface markers for human ABC, they were not as robust as scRNA-seq analysis and cannot capture the general ABC population.

To determine what may be the key TFs required for the formation of ABCs we examined which TFs were significantly upregulated in ABCs compared to cMBCs within our dataset. This revealed the top 5 candidate TFs: SOX5; ZBED2; BHLHE40; ZEB2; and ZBTB32 (**Figure 1G and H**). Ideally, the key TFs should promote ABC genes and suppresses cMBC genes, therefore, we performed an analysis to determine which TF correlated most strongly in individual cells with the expression of the ABC geneset, the surface expression of CD11c, and which were most inversely correlated with cMBC gene expression and surface CD27 expression. In this analysis ZEB2 was the top candidate, though expression of BHLHE40 and SOX5 also correlated with the expression of ABC genes (**Figure 1I**).

### Murine ABCs can be generated in vitro and after sporozoite immunization in vivo

To investigate the role of individual TFs we sought to develop experimentally tractable models of ABC formation. In agreement with others (27), we identified a population of CD11c^+^ B cells after infection with rodent malaria *P. chabaudi* (**Figure S2A and B**). RNA-seq analysis revealed that these cells had a distinct transcriptional profile from cMBCs (**Figure S2C**) characterized by the expression of many genes homologous to those expressed by human ABCs (**Figure S2D**). However, as ABCs are clearly prominent in the blood and tonsils of otherwise healthy donors we also wanted to establish a model of ABC formation in a non-disease state. Our previous analysis showed that immunization of humans with *Plasmodium* sporozoites induces a population of ABCs specific for the immunodominant *P. falciparum* circumsporozoite protein (PfCSP) even after primary immunization (8). We therefore asked if the same might be true in mice. To assess this we transferred PfCSP-specific Igh^g2A10^ cells (28) to MD4 recipients and immunized with *P. berghei*-PfCSP sporozoites (PfCSP-SPZ) that express PfCSP in place of the endogenous *P. berghei* CSP protein (29). The use of recipient Ig-transgenic mice which express an irrelevant BCR - in this case specific for hen egg lysozyme - is an established approach for the study of rare memory cell populations (30). Immune responses were assessed 4, 10 and 21 days post-immunization (**Figure 2A**). At all time points CD11c^+^ and CD11c^−^ switched B cells were identified in the spleen, LN, blood and bone marrow of mice (**Figure 2B and C**), with ABCs being most enriched in the spleen where they peaked on day 10 accounting for ~15% of memory Igh^g2A10^ cells (**Figure 1C**). The identity of these cells as ABCs was confirmed by bulk RNA-seq analysis of CD11c^+^ and CD11c^−^ cells at each timepoint. CD11c^+^ B cells were clearly separated from CD11c^−^ B cells by PCA analysis especially at days 10 and 21 (**Figure 2D**) and they also up-regulated many known ABC genes and down-regulated cMBC genes (**Figure 2E**).

**Figure 2.**
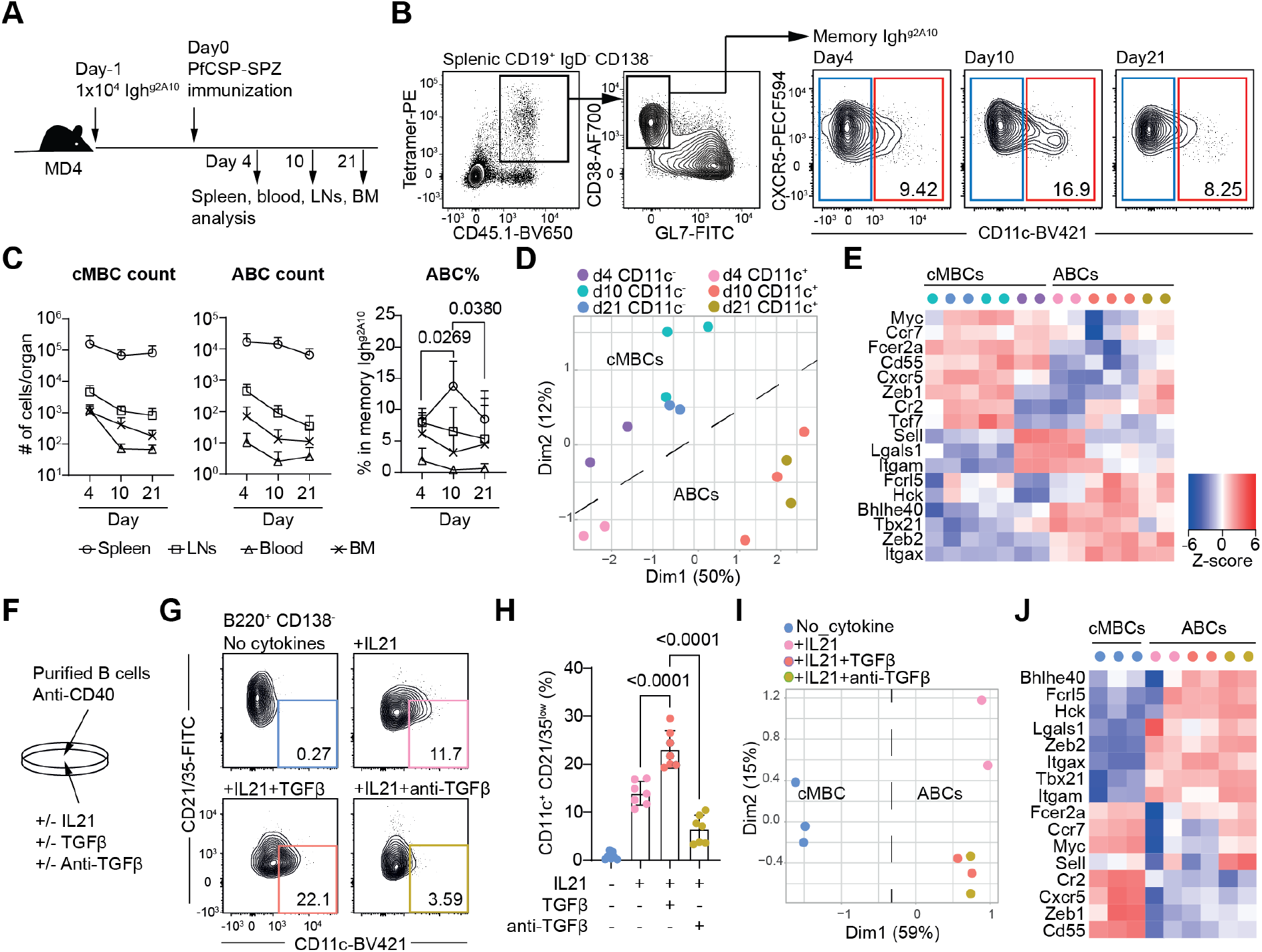
Murine ABCs can be generated in vitro and after sporozoite immunization in vivo. (A-E) 10^4^ Igh^g2A10^ B cells were IV injected into MD4 mice, followed by PfCSP-SPZ immunization. Spleens, blood, LNs and BM were analysed on day4, day10 and day21 after immunization. Splenic memory Igh^g2A10^ CD11c^−^ and CD11c^+^ cells were also sorted for bulk RNA-seq analysis. (A) Experiment design. (B) Representative FACS plot for splenic ABCs percentages in memory Igh^g2A10^. (C) Statistics of cMBC and ABC Igh^g2A10^ total cell count and percentage in memory cells. (D) PCA plot comparing the transcriptomes of CD11c^−^ and CD11c^+^ memory Igh^g2A10^ cells on all timepoints. (E) Heatmap showing the expression of ABC and cMBC signature genes in all samples. (F-J) MACS enriched mouse B cells were cultured with anti-CD40 w/wo IL-21 and TGFβ for 4 days followed by FACS analysis or sorting (B220^+^ CD138^−^) for bulk RNA-seq analysis. (F) Experiment design. (G) Representative FACS plot and (H) statistics of CD11c^+^ CD21/35^lo^ cells for each culture condition. (I) PCA plot comparing the transcriptomes of B cells cultured under indicated conditions. (J) Heatmap showing the expression of ABC and cMBC signature genes in all samples. The results in (C) and (H) were pooled from two independent experiments. The P values were calculated by One-way ANOVA.

To further aid in studies of ABC development we also attempted to define conditions to induce ABCs in vitro (**Figure 2F**). In our hands we found that IL21 was critical for driving the development of a CD11c^+^ CD21^−^ cells in vitro after stimulation with anti-CD40 (**Figure 2G-H**), consistent with previous reports about the essential role of IL21 in ABC development (20, 31–33). Based on our analysis of tonsil single cell dataset we further found that ABCs expressed many genes in the TGFβ signalling pathway (**Figure S2E-F**). In agreement with this, the addition of TGFβ modestly enhanced ABC formation, while anti-TGFβ almost completely abolished ABC formation in vitro (**Figure 2G-H**). TGFβ blockade in vivo also mildly impaired ABC formation upon PfCSP-SPZ immunization (**Figure S2G-I**). However, we found the presence of TGFβ or not in the culture did not affect the general ABC programming as illustrated by RNA-seq analysis (**Figure 2I-J**), suggesting TGFβ may play a redundant role during ABC formation.

We then asked whether ABCs originated from different contexts share a similar global transcriptomic profile. We performed GSEA across different RNA-seq datasets (including our in vitro and in vivo mouse datasets, human tonsil dataset and datasets from previous publications) and found that these ABCs were transcriptomically highly similar (**Figure S2J**). Such resemblance was likely controlled by specific transcription factors (TFs), therefore we further analyzed the differently expressed TFs in our in vitro and in vivo mouse models. Consistent with our results in human tonsils, we found that Zeb2 was again highly upregulated in ABCs in both mouse datasets (**Figure S2K**).

### A CRISPR/Cas9 knock-out screen identifies Zeb2 is required for ABC formation

To help systemically identify the key TFs for ABCs, we developed an optimized Cas9 ribonuclear protein (RNP) mediated gene knock-out (KO) method to delete individual TFs in cultured B cells. The efficient delivery of RNP by nucleofection has previously been shown to require B cell pre-activation (34, 35). In agreement with this we found anti-CD40 pre-activation was optimal for transfection (**Figure S3A-C**). To further optimize the gene KO efficiency, we co-transfected our target gene RNP with a sub-optimal concentration of GFP RNP into GFP^+^ B cells. We reasoned that successfully transfected cells would be GFP^−^ by flow cytometry (**Figure S3D**). We validated this method by KO of Bcl6, the master TF for GC B cells and as predicted, found an enhanced KO efficiency in GFP^−^ cells (80-90% indel) compared to GFP^+^ counterparts (60-75% indel) as determined by Sanger sequencing (**Figure S3E**). The robustness of this method was further confirmed by the nearly complete abolishment of GC B cells in the transferred Bcl6KO GFP^−^ Igh^g2A10^ cells following PfCSP-SPZ immunization (**Figure S3F-H**). Because previous studies have been conflicting on whether ABC formation requires GC (20, 26), we further analyzed the percentages of ABCs in memory Igh^g2A10^ B cells and found Bcl6 was not required for ABC formation (**Figure S3G, S3I**). Interestingly, we found GFP KO mildly but significantly affects ABC formation (**Figure S3G, S3I**). Therefore, in the following experiments we only compared ABC% in GFP^−^ cells to avoid the confounding effects of GFP expression.

Because ABCs are a highly conserved population (**Figure S2J**), we hypothesized that the key TFs for ABC should be shared by different RNA-seq datasets. Therefore, we calculated the upregulated TFs in each of our RNA-seq datasets and identified 5 highly shared TFs including Zeb2, Tbx21, Bhlhe40, Tcf7l2 and Mafb (**Figure 3A**). We also analyzed Bhlhe41 and Maf because they belong the same families as Bhlhe40 and Mafb respectively. Runx2 was previously proposed to play a role in human ABC formation (36) and so was included in our screen. We then used our optimized Cas9 KO method to study the roles of these TFs. We subsequently measured ABC formation either in vitro or by transfer of the Cas9 RNP transfected Igh^g2A10^ cells into MD4 mice with subsequent PfCSP-SPZ immunization for assessment of in vivo ABC formation (**Figure 3B**). As expected, the KO of Il21r resulted in reduced ABC formation compared to nontarget control by in vitro culture (**Figure 3C-D**) and a trend of reduction by in vivo PfCSP-SPZ immunization model (**Figure 3E-F**). Notably, in this system only the KO of Zeb2 reduced ABC formation in both our in vitro and in vivo models (**Figure 3C-F**). Therefore, we concluded that Zeb2 is likely a key TF for ABC formation.

**Figure 3.**
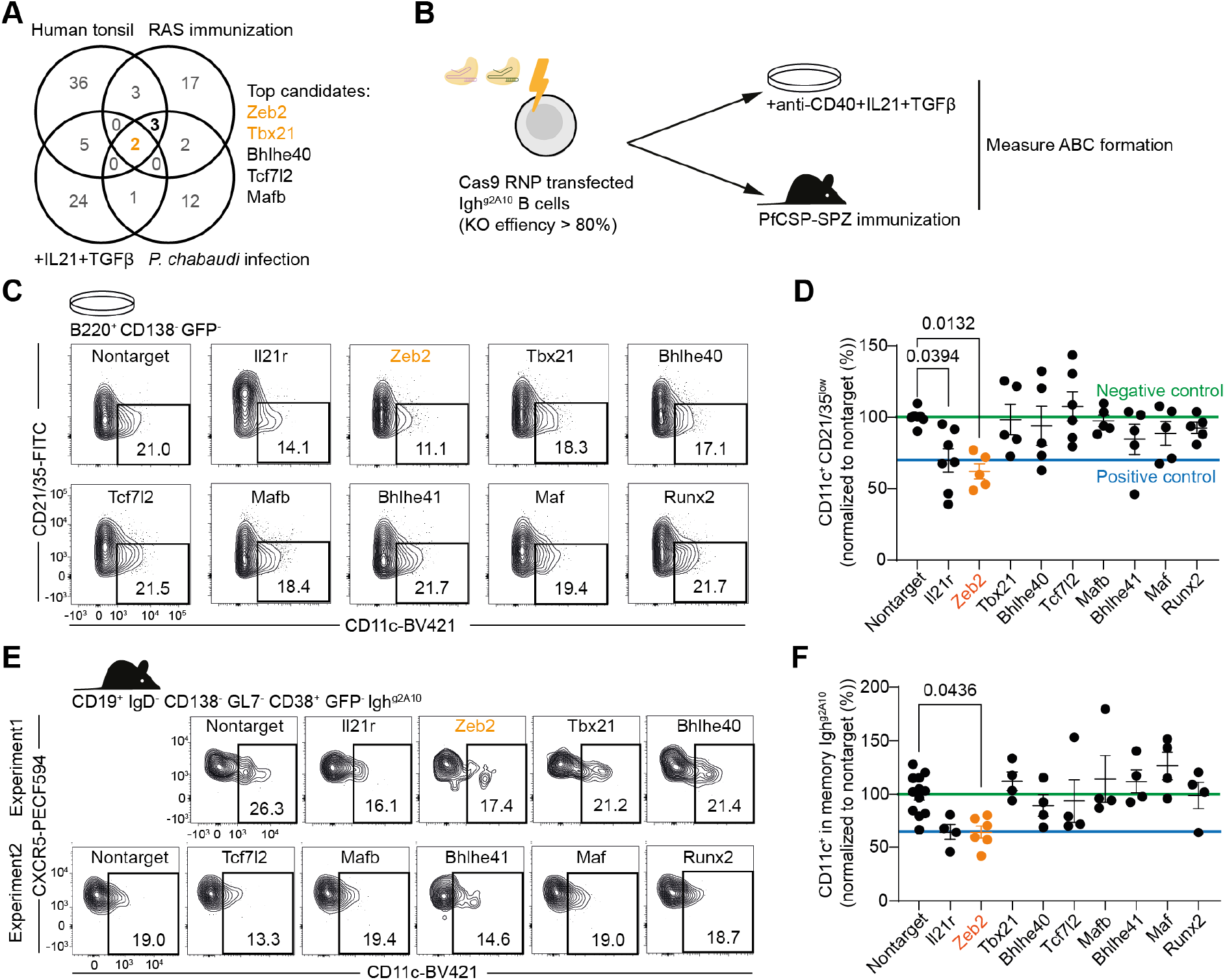
A CRISPR/Cas9 knock-out screen identifies Zeb2 is required for ABC formation. (A) Venn diagram showing the numbers of significantly upregulated TFs in indicated RNA-seq datasets by comparing ABC vs cMBC. Threshold were set as follows: Human tonsil (FDR ≤0.01), the other datasets: (P ≤0.05, Log2FC ≥0.5). (B-F) MACS purified Igh^g2A10^ ^B^ cells were pre-activated by 10ug/ml anti-CD40 for 24h, followed by Cas9-RNP nucleofection and a proportion of cells were culture for 4 days with anti-CD40, IL-21 and TGFβ followed by FACS analysis. The rest of cells were transferred into MD4 followed by PfCSP-SPZ immunization. (B) Experiment design. (C) Representative FACS plots and (D) statistics showing the ratio of CD11c^+^ CD21/35^lo^ cells in CD138^−^ B cells transfected by different Cas9 RNPs. (E) Representative FACS plots and (F) statistics showing the percentages of ABCs in GFP^−^ memory Igh^g2A10^ B cells. The results were pooled from four independent experiments for (D) and (F). Each Cas9 RNP was tested by at least two independent experiments with two to three replicates for each experiment. Data normalization was performed by calculating the percentage of nontarget in each individual experiment and then the data was pooled. The P values were calculated by One-way ANOVA.

### Zeb2 deficiency selectively impairs the development of ABCs in mice and humans

Because our Cas9 system did not fully disrupt the *Zeb2* locus (**Figure S4A**), and the pre-activation by anti-CD40 might skew the normal development of other B cell lineages like PB/PC and GC B cells, we further studied the role of Zeb2 in ABC formation using a Cre-lox system. Given that Zeb2 is required for B cell development during pre-pro to pro B cell transition (37, 38), we crossed previously described *Zeb2^fl/fl^* mice (39) with a *Cd23^Cre/+^* line such that *Zeb2* would only be ablated in mature B cells. Control mice were *Cd23^Cre/+^* and either heterozygous or negative for the *Zeb2^fl^* allele. In vitro differentiation experiments with these cells reproduced our previous finding and again, we found a strong reduction in ABC formation in both heterozygous *Zeb2^fl/+^* and *Zeb2^fl/fl^* B cells compared to *Cd23^Cre/+^* control B cells (**Figure S4B-D**). We also checked the expression of other ABC markers and again found an overall impairment of the up-regulation of ABC markers CD11b, CD72, CD19 and the down-regulation of cMBC markers CD21, CD23, CD55 (**Figure S4E-F**), suggesting that Zeb2 is required for the acquisition of the general ABC phenotype.

To investigate whether Zeb2 was critical for ABC formation in vivo we further crossed *Cd23^Cre+^ Zeb2^fl/fl^* mice to the *Igh^g2A10^* background and transferred splenocytes from the resulting mice into MD4 animals to investigate the outcome of B cell specific Zeb2 deficiency in response upon PfCSP-SPZ immunization (**Figure 4A**). We found Zeb2 deficient *Igh^g2A10^* cells had significantly reduced ABC responses with a lower percentage of ABCs compared to mice that received *Zeb2^+/+^ Cd23^Cre/+^ Igh^g2A10^* B cells with *Zeb2^fl/+^Cd23^Cre/+^ Igh^g2A10^* heterozygotes having an intermediate phenotype (**Figure 4B-D**). Of note however the numbers and proportions of other B cell populations including total IgD^−^ cells, PB/PC, GC B cells and cMBCs were not affected (**Figure 4B-C**), suggesting Zeb2 deficiency selectively impairs ABC formation without affecting other B cell populations. Interestingly, this result also suggests ABCs do not play a significant role in the primary B cell response, though we cannot exclude a role in the development of recall responses.

**Figure 4.**
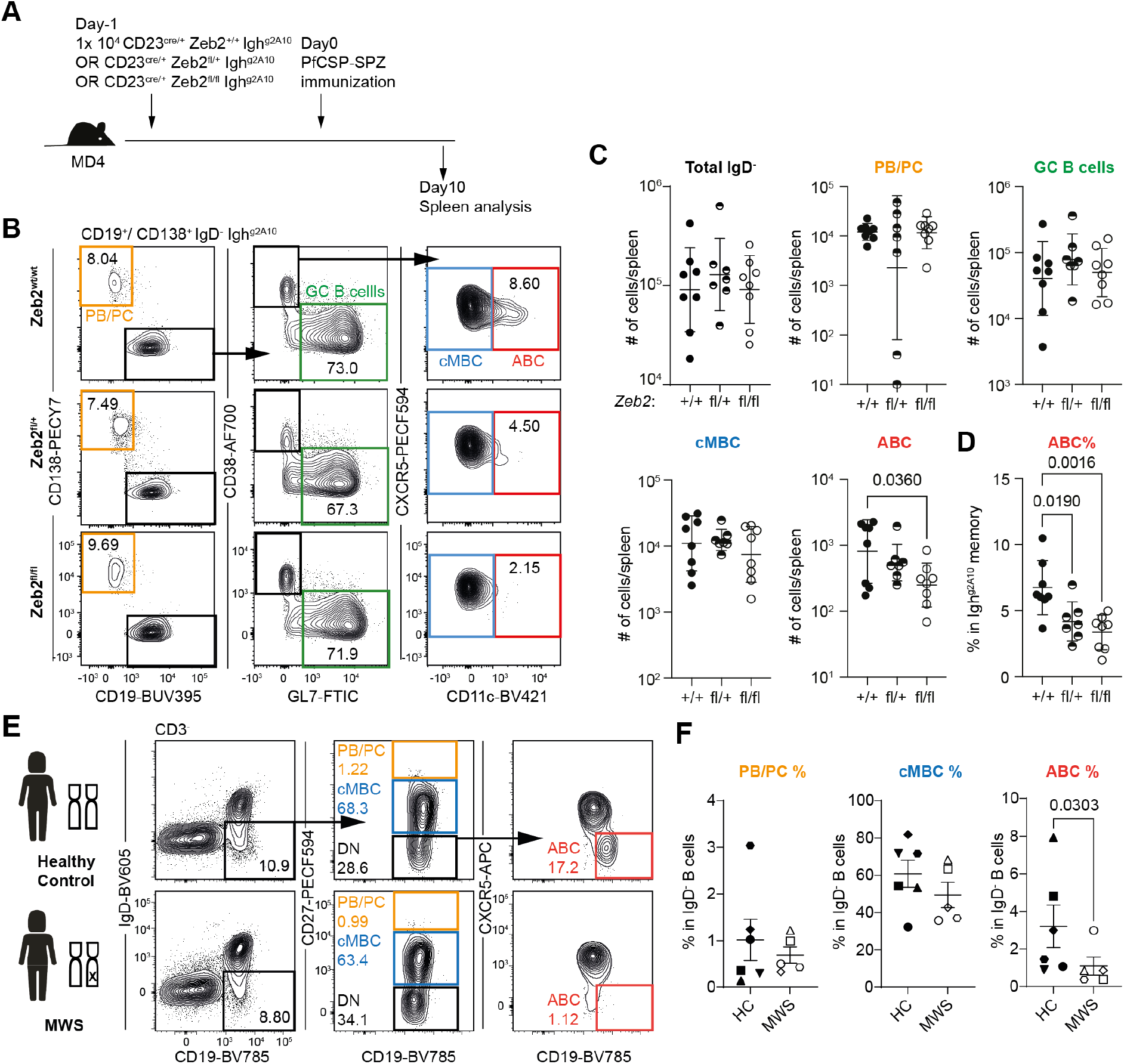
Zeb2 deficiency selectively impairs the development of ABCs in mice and humans. (A-D) 10^4^ Cd23^cre/+^ Zeb2^+/+, fl/wt^ ^or^ ^fl/fl^ Igh^g2A10^ B cells were transferred into MD4 mice followed by PfCSP-SPZ immunization, the spleens were analyzed on day10. (A) Experiment design. (B) Representative FACS plot and (C, D) statistics showing the formation of indicated Igh^g2A10^ B cell populations. (E-F) PBMC samples from 5 MWS patients and 6 healthy controls were FACS analyzed. (E) Representative FACS plots and (F) statistics showing the percentages of indicated B cell populations. The results for (C, D) were pooled from two independent experiments. The P values were calculated by one-way ANNOVA for (D) and Mann Whitney U test for (F).

Finally, to determine whether ZEB2 may also play a role in ABC development in humans we examined B cells in patients with Mowat-Wilson Syndrome (MWS). MWS is a rare genetic disease in humans (incidence: 1 per 50000-70000 live births (40)) caused by ZEB2 haploinsufficiency resulting in developmental abnormalities. Overt immune system defects have not been reported in MWS patients and they have normal immunoglobulin levels and responses to vaccinations (41). However, previous studies also reported the MWS patients have significantly reduced percentages of CD8^+^ T cell in lymphocytes (41) and decreased percentages of B cells in lymphocytes though this was not statistically significant (42). The fact that murine heterozygous *CD23^Cre/+^ Zeb2^fl/+^* cells had reduced ABC formation both in vitro and in vivo implied that it might be possible to observe a similar phenotype in MWS patients. We therefore analyzed the PBMC samples from 5 MWS patients and 6 age/sex matched healthy controls (HCs) and calculated the percentages of ABCs as well as PB/PC and cMBCs in IgD^−^ B cells. We analyzed the percentage of ABCs in IgD^−^ B cells because different people have large variation on the numbers of IgD^+^ naïve B cells due to different antigen exposure history. MWS patients did not have decreased B cell percentages among lymphocytes suggesting they have normal B cell development (**Figure S4G**). However, MWS patients had significantly decreased percentages of ABCs in IgD^−^ B cells compared to HCs while the development of other B cell subsets like PB/PC and cMBCs appeared to be normal (**Figure 4E-F**). Consistent mouse and human results suggest Zeb2 deficiency selectively impairs ABC formation, further strengthening our conclusion that Zeb2 as a key TF for ABC formation.

The identification of ABCs, not only in disease states but also in normal responses raises key questions about their functional role. In SLE they have been proposed to be a pathogenic sub-type responsible for the secretion of autoantibodies (10, 12, 43). In this context a recent pre-print study also suggests that Zeb2 may be the a key driver of the ABC program by binding to the *Itgax* and *Itgam* TF regulatory locus and activating via the Jak-Stat pathway (44). Another possible function for ABC is antigen presentation to T cells (43, 45) especially membrane bound antigen (46). Consistent with this, Zeb2 has been shown to be critical for the development of cDC2 cells responsible for the priming of Th2 cells upon *Heligmosomoides polygyrus* infection (47, 48).

Here we show that Zeb2 is selectively required for the ABC formation that is observed after vaccination in mice and natural antigen exposure in humans. However, a key limitation in our study is we have been unable to detect any obvious impairment of the B cell responses in Zeb2 deficient mice and humans that would otherwise be indicative of the normal function of these cells. Moreover, our current knowledge of ABC biology is hampered by the lack of a valid method to specifically deplete ABCs for functionally studies. Therefore, the *CD23^cre/+^ Zeb2^fl^* mice may be a useful tool for this purpose. A second limitation is that we have been unable to overexpress Zeb2 in B cells to determine if expression of this transcription is sufficient to drive the ABC transcriptional program. Nonetheless, the finding of the key TF that selectively required for ABC formation is conceptually important because it demonstrates that these alternative B cells represent a distinct lineage. Our future studies will extend from the context of primary immunization to investigate the responses to antigen recall and to fulminant malaria infection in Zeb2 deficient animals.

## Materials and Methods

### RESOURCE AVAILABILITY

#### Lead contact

Further information and requests for resources and reagents should be directed to and will be fulfilled by the lead contact, Ian A. Cockburn.

#### Materials availability

This study did not generate new unique reagents.

#### Data and code availability

The original and processed single-cell and bulk RNA-seq data have been deposited at GEO and will be publicly available as of the date of final publication. The original code for bioinformatic analysis is available in this paper’s supplemental information. Any additional information required to reanalyze the data reported in this paper is available from the lead contact upon request.

### EXPERIMENTAL MODEL AND SUBJECT DETAILS

#### Ethics statement

The study on human tonsillar cells was approved by the Australian National University Human Experimentation Ethics Committee and Australian Capital Territory Government Centre for Health and Medical Research (Protocol numbers: ANU 2012/517; ACT Health ETH.4.06.268). The study on MWS patient and age/gender matched healthy controls was approved by the Sydney Children’s Hospitals Network Human Research Ethics Committee (MWS patients, protocol number: LNR/15/SCHN/195) and St Vincent’s Hospital Sydney Human Research Ethics Committee (healthy controls, protocol numbers: HREC/13/SVH/145; 2019/ETH03336). All research involving human samples was conducted in accordance National Statement on Ethical Conduct in Human Research 2018. All animal procedures were approved by the Animal Experimentation Ethics Committee of the Australian National University (Protocol number: 2019/36). All research involving animals was conducted in accordance with the National Health and Medical Research Council’s Australian Code for the Care and Use of Animals for Scientific Purposes and the Australian Capital Territory Animal Welfare Act 1992.

#### Human samples

Human tonsils were obtained from two children (median age (range): 4.5 (4-5) yrs, 2/2 female) underwent tonsillectomy. PBMC samples were collected from five MWS patients (median age (range)= 12 (3-21) yrs, 1/6 female) and six healthy controls (median age (range)= 21.5 (21-24) yrs, 2/6 female).

#### Mice

C57BL/6 mice, MD4 (49) and Igh^g2A10^ (28) were bred in-house at the Australian National University. All mice were on a C57BL/6 background. Mice used for the experiment were 5 to 8 weeks, and they were age matched for each experiment groups. Mostly female mice were used throughout the experiments, and were bred and maintained under specific pathogen free conditions in individually ventilated cages at the Australian National University.

#### Parasites

*P. berghei* parasites engineered to express *P. falciparum* CSP in place of the endogenous *P. berghei* CSP molecule were used throughout the study for immunization (29) and were maintained by serial passage through *Anopheles stephensi* mosquitoes. *P. chabaudi* parasites were maintained by serial passage through mouse blood stage infection.

### METHOD DETAILS

#### Tonsiliar cell preparation for scRNA-seq

Tonsils were cut into fine pieces and smashed against a 70μm cell strainer. The tonsillar mononuclear cells were then purified by Ficoll-Paque Plus (GE) density gradient separation. After Fc blocking, tonsillar mononuclear cells were incubated in antibody cocktail on ice for 30 min followed by flow cytometry sorting for total B cells, GC B cells and memory B cells. The sorted cells were then incubated with Totalseq-C antibodies (1:50 dilution) on ice for 30 min, washed 3 times (by default, top up with 2% FBS (Gibco) in DPBS (Sigma) to 10 mL and centrifuging at 800g for 3-5min), counted and mixed at 1:1:1 ratio. 1 × 10^4^ mixed cells were loaded onto each lane of the 10X Chromium platform (10X Genomics). Library preparation was completed by Biomedical Research Facility (BRF) at the JCSMR following the recommended protocols. Libraries were sequenced using the NovaSeq6000 (Illumina). The 10X Cell Ranger package was used to process transcript, CITE-seq and VDJ libraries for downstream analysis. Details of all key reagents for single-cell RNA-seq are given in the Key resources table.

#### Immunization and infection

For immunization, mice were IV injected with 5 × 10^4^ irradiated (15kRad) Pb-PfSPZ in DPBSnbsdissected by hand from the salivary glands of *Anopheles stephensi* mosquitoes. For infection, mice were IV injected with 1 × 10^5^ *P. chabaudi* infected red blood cells in DPBS. 100ug anti-TGFb antibody (Bioxcell) per mouse per dose was administered by IP injection.

#### Cas9 RNP preparation and nucleofection

Anti-PE MACS beads (Miltenyi) enriched B220-PE^+^ GFP^+^ splenic mouse B cells were resuspended in complete RPMI media (Gibco) at 5 million cells per mL and stimulated by 10ug/mL anti-CD40 antibody for 24h. After wash once by DPBS, 1 million pre-activated B cells were resuspended in RNP solution for each nucleofection. RNP solution (for each nucleofection) was prepared by mixing 250 nmol target gene sgRNA+ 50 nmol eGFP sgRNA+ 50 nmol SpCas9 2NLS Nuclease (Synthego) in 50 μL mouse B cell nucleofection buffer (Lonza) at RT for 30min. The nucleofection was then performed by AMAXA Nucleofector I device X-01 program (Lonza). The nucleofected cells were rested in the cuvette at RT for 10min before being transferred out for downstream applications. Details of all reagents for Cas9 RNP preparation are given in the Key resources table.

#### Cell culture

MACS enriched naïve mouse B cells or Cas9 RNP transfected B cells were cultured for 4 days in 200 μL complete RPMI media per well of 96-well plate (Nunc) in the presence of 3 μg/ml anti-CD40 antibody (Biolegend) w/wo 50 ng/mL IL-21, 5ng/mL human TGFb (Peprotech) and 10ug/mL anti-TGFb. Media is changed on day2 by gently aspirating 180 μL old media and adding 180 μL new media (we found changing media, especially the renewal of IL-21, is essential for successful ABC differentiation).

#### Flow Cytometry

For phenotyping MWS patients and healthy controls, 100 μL of fresh Na heparin anti-coagulated peripheral blood was stained with monoclonal antibodies (mAb) according to the manufacturers’ directions, incubated for 15 min at RT, lysed with Optilyse C (Beckman Coulter) for 10 min at RT, washed and resuspended in 250 μL 0.5% paraformaldehyde in PBS for further flow cytometry analysis. For mouse studies, ACK lysis buffer treated splenic cells, blood cells, bone marrow cells or cultured B cells were Fc blocked on ice for 10min and top up by the same volume of 2× concentrated antibody cocktail for another 1h on ice, followed by one wash and flow cytometry analysis. Details of antibodies are given in the Key resources table.

#### Sanger sequencing to calculate indel frequency

Genomic DNA were extracted from untreated or Cas9 RNP nucleofected cells (24h after transfection) according to manufacturer’s instruction (Monarch). The ~500bp DNA sequence flanking the expected cutting site were amplified by PCR and Sanger sequenced using another inner primer. The unedited and Cas9 edited sequences were then analyzed by the Synthego ICE tool for indel frequency calculation. Primer sequences to amplify the Bcl6 locus: forward: TTTGGGCCTTACAGGGCAAG; reverse: GCTTGTTTTACAGCGGCCTG; inner: GTGTCTGCCTAGAGCATGTGA. Primer sequences to amplify the Zeb2 locus: forward: TCCAGAAGGGCGATGGAAAG; reverse: TGAATATGTGGGGGCATTGGT; inner: CACTTACCTGGACCGGCTAC.

#### RNA extraction and bulk RNA-seq

2 × 10^3^ to 1 × 10^5^ sorted B cells per sample were centrifuged at 800g for 8min and total RNA was extracted using RNeasy Micro Kit (Qiagen) following the recommended protocol. The RNA concentration and RNA integrity number (RIN) were measured by Bioanalyzer RNA 6000 pico assay (Agilent). Total RNA samples with RIN ≥6.0 and concentration ≥30pmol/μL were submitted to BRF at the JCSMR for downstream process. Library preparation and sequencing was completed using NEBNext® Single Cell/Low Input RNA Library Prep Kit (NEB) and NextSeq 550 (Illumina) following recommended protocol.

#### Bioinformatic analysis of scRNA-seq dataset

The analysis of 10X scRNA-seq dataset has been described in our previous publication (8). In brief, transcripts and CITE-seq library outputs were loaded into Seurat package (50) for unwanted cell removal, clustering, annotation and visualization such as UMAP, violin plot, feature plot and dot plot. VDJ analysis were performed by IMGT/HighV-QUEST for SHM frequency and Ig isotype assignment. The FACS plots were made by Flowjo using the exported CITE-seq antibody expression values. All the details can be found in the original code which is available in this paper’s supplemental information.

#### Bioinformatic analysis of bulk RNA-seq dataset

The bulk RNA-seq dataset were analyzed by published pipelines (51, 52). The generated single-end 75bp reads were processed under the Galaxy project online platform and the mapped count files were then analyzed by Limma for cpm normalization, PCA plot generation and differentially expressed genes calculation. Gene set enrichment analysis (GSEA) was performed by fgsea. The details of pre-ranked genes and gene sets are available in this paper’s supplemental information.

### QUANTIFICATION AND STATISTICAL ANALYSIS

Statistical analysis was performed by Prism software (GraphPad). All data were assumed Gaussian distributed thus comparisons between two groups were performed by two-tailed parametric (for mice data) or nonparametric (for human data) t-test and multiple comparisons were performed by one-way ANOVA post hoc Dunnett or Tukey test using the default settings of Prism. P values < 0.05 were considered significant and specified in the paper.

## Acknowledgments

We thank supports from Harpreet Vohra and Michael Devoy of the Imaging and Cytometry Facility at the Australian National University.

## Author contributions

Study design: X.G., K.F., J.J.Z. and I.A.C..; Generation of human tonsil scRNA-seq dataset: Q.S., X.G., M.N. and M.C.C.; Bioinformatic analysis of scRNA-seq dataset: X.G. and J.A.R.; Generation of mouse bulk RNA-seq datasets: X.G., B.D. and M.N.; Bioinformatic analysis of bulk RNA-seq datasets: X.G., B.D. and R.J.; CRISPR-Cas9 system optimization: X.G. and J.H.; Mouse experiments: X.G.; MWS patients and healthy donors sample collection and flow cytometry analysis: K.F.; C.M.M.; J.Z.; X.G.; and I.A.C.; Manuscript writing: I.A.C. and X.G. All authors reviewed and approved the final version.

## Supplementary Figures

**Figure S1.**
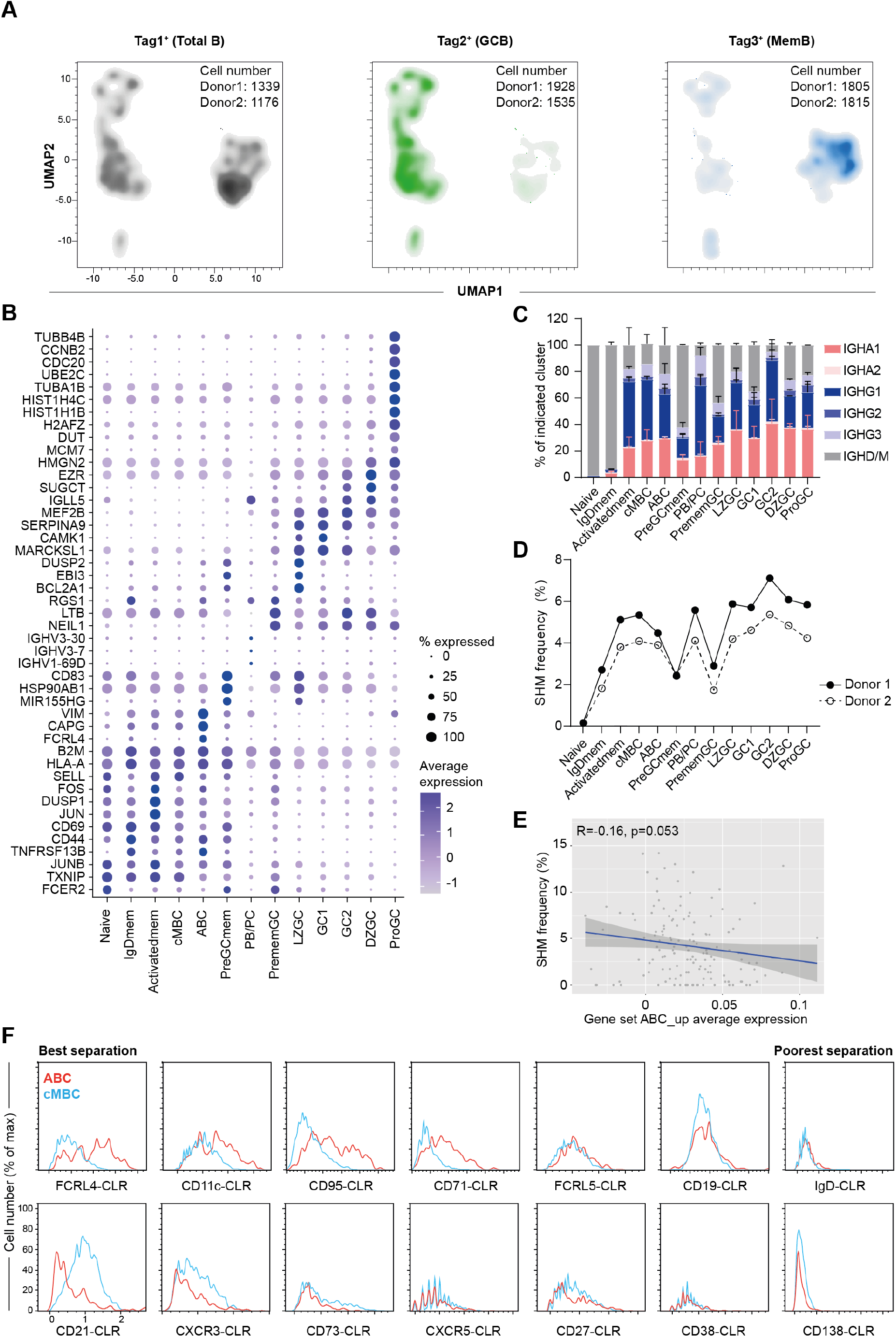
scRNA-seq analysis on human tonsil B cells. (A) UMAP of sorted total B cells, GC B cells and memory B cells labelled by Hashtag 1, 2 and 3 respectively.(B) Dot plot showing the expression of marker genes for each cell cluster identified by FindAllMarkers function. (C-E) VDJ sequences were mapped to IMGT database through V-QUEST to identify heavy chain usage and mutation frequency for each cell. (C) The distribution of immunoglobulin isotypes for each cell clusters. (D) Statistics of SHM frequency for each cell clusters. I Pearson correlation analysis between the SHM frequency and the average expressions of ABC genes in ABC cluster. (F) The histograms comparing the expressions of all markers between ABC and cMBC identified by CITE-seq.

**Figure S2.**
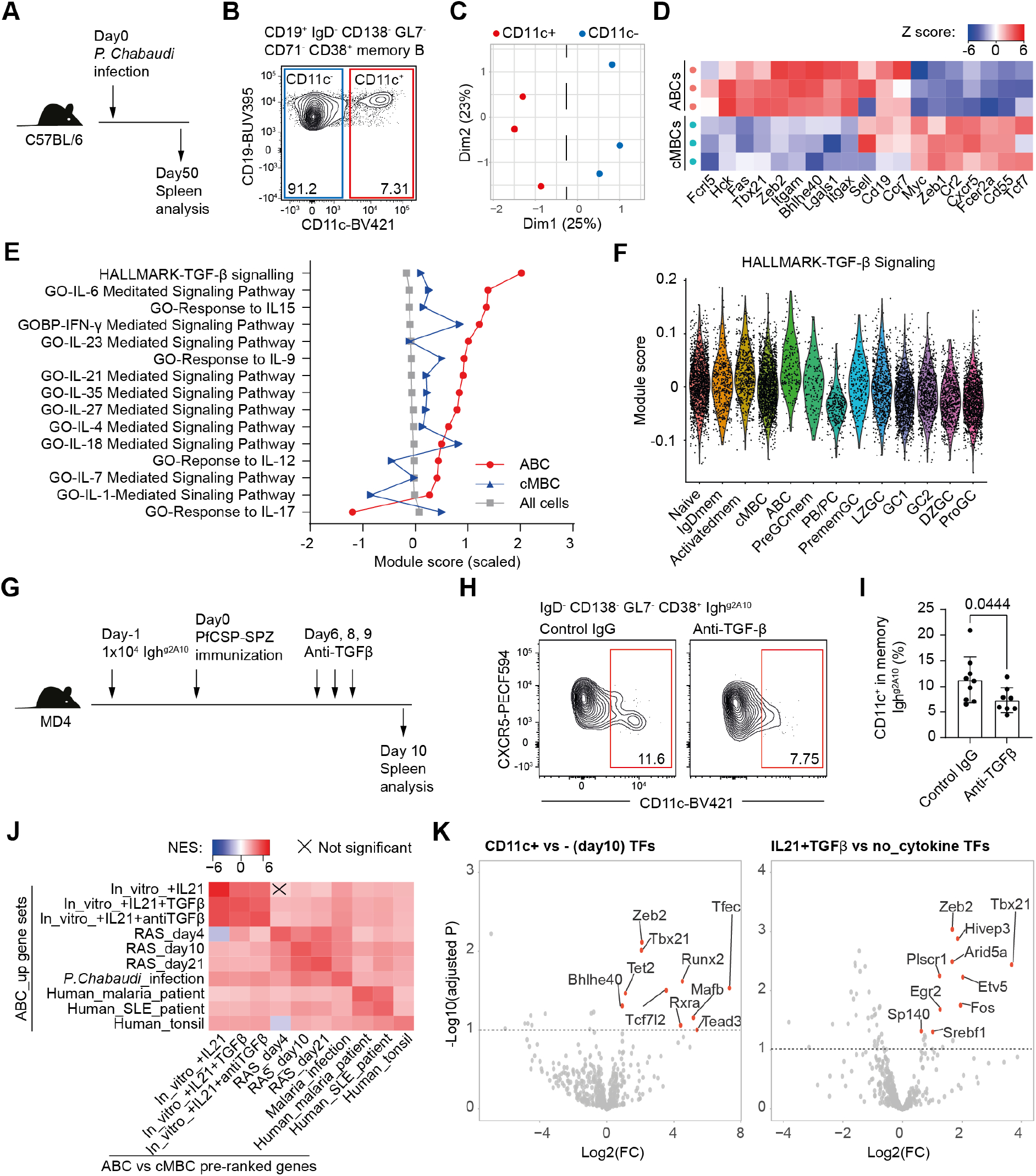
Murine models of ABC formation. (A-D) WT mice were infected by IV injecting 10^5^ P. chabaudi iRBC and spleens were analysed on day50. CD71^−^ resting CD11c^+^ and CD11c^−^ memory B cells were also sorted for bulk RNA-seq analysis. (A) Experiment design. (B) Representative FACS plot for ABC formation. (C) PCA comparing the transcriptomes of CD11c^+^ and CD11c^−^ cells. (D) Heatmap showing the expression of ABC and cMBC signature genes in all sampleI(E) Average enrichment scores for cytokine pathways were calculated and summarized and ranked based on the values for ABC cluster. (F) The violin plot showing the enrichment scores for TGFβ signalling pathway in all clusters. (G-I) 10^4^ Igh^g2A10^ B cells were transferred into MD4 mice followed by PfCSP-SPZ immunization. The mice were IP injected with either control IgG or anti-TGFβ on day6, 8 and 9. The spleens were analysed on day10. (G) Experiment design, (H) representative FACS plots and (I) statistics of CD11c^+^ ABC % in Igh^g2A10^ cells. (J) ABC vs cMBC signature genes and pre-ranked gene lists were calculated based on RNA-seq datasets in this study and other publications (10, 25). Cross-dataset GSEA were performed and the normalized enrichment scores (NES) and significance were summarized as heatmap. Human gene symbols were transformed into mouse homologous genes for this purpose. RAS: PfCSP-SPZ immunization. (K) The volcano plots showing the top10 upregulated TFs in ABCs derived from in vitro culture and PfCSP-SPZ immunization. Result for (I) was pooled from two independent experiments. The P value was calculated by student’s t-test

**Figure S3.**
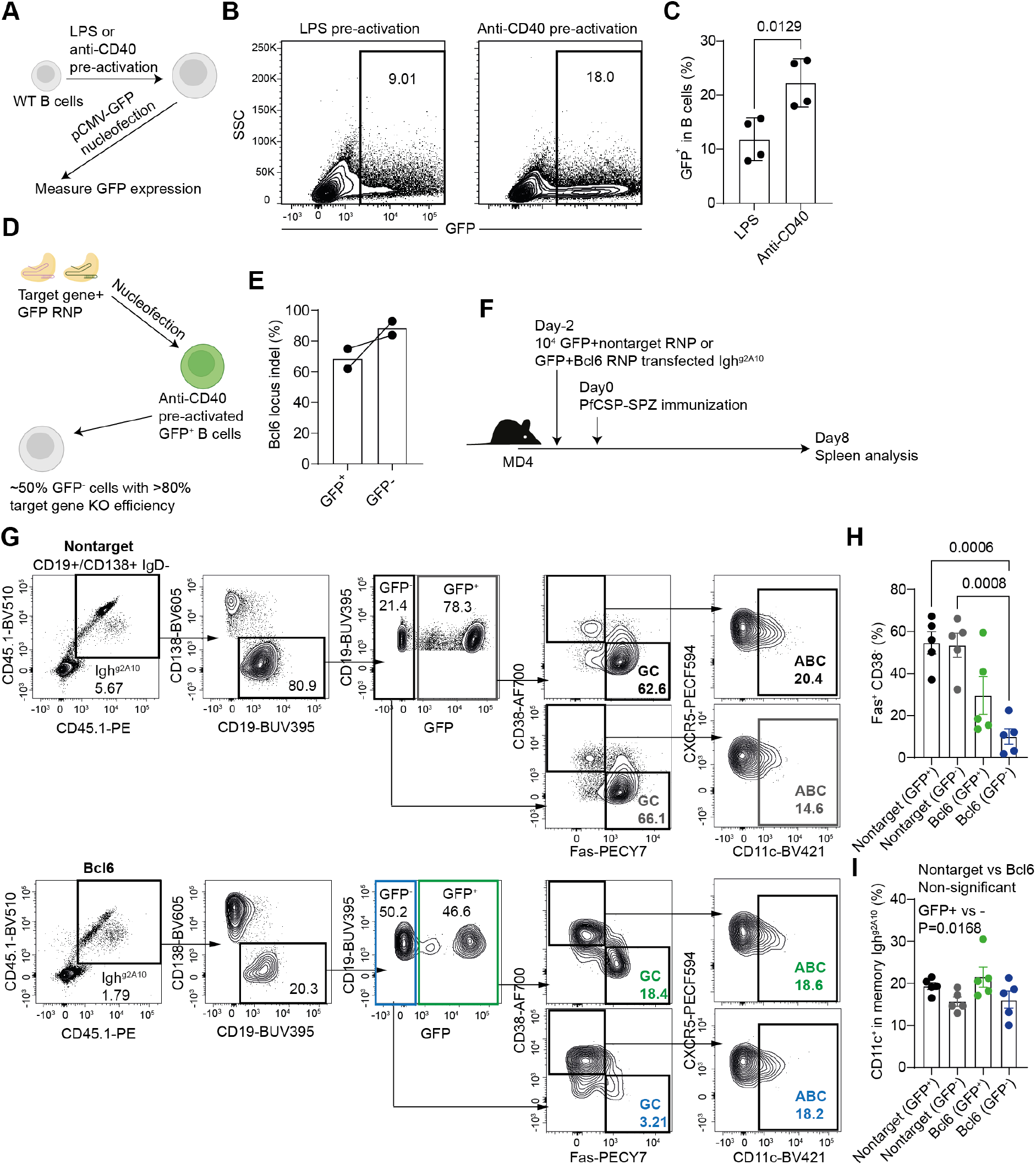
An optimized Cas9-RNP based gene knock-out method in mouse B cells. (A-C) MACS enriched mouse B cells were pre-acitvated by 10ug/ml LPS or anti-CD40 for 24h, followed by nucleofection with 1ug Lonza pCMV-GFP plasmid. The expressions of GFP were measured 24h after the transfection. (A) Experiment design. (B) Representative FACS plots and (C) statisitcs showing the expression of GFP in transfected B cells. (D-I) MACS enriched GFP-expressing Igh^g2A10^ B cells were pre-activated by anti-CD40, nucleofected with Bcl6 and GFP Cas9 RNPs and transferred into MD4 mice followed by PfCSP-SPZ immunization. A small aliquot of cells was rested in anti-CD40 supplemented media for 24h, followed by gDNA extraction, PCR and Sanger sequencing to evaluate indel frequency by ICE analysis. (D) Experiment design. (E) Statistics comparing the Bcl6 indel frequency in GFP^+^ and GFP^−^ transfected cells. (F) Experiment design, (G) representative FACS plots and statistics showing the percentages of (H) Fas^+^ CD38^−^ GC B cells and (I) ABCs in transfected memory Igh^g2A10^ B cells. Results were pooled from two independent experiments for (C, E, H, I). The P values were calculated by student’s t test for (C), one-way ANNOVA for (H) and two-way ANNOVA for (I).

**Figure S4.**
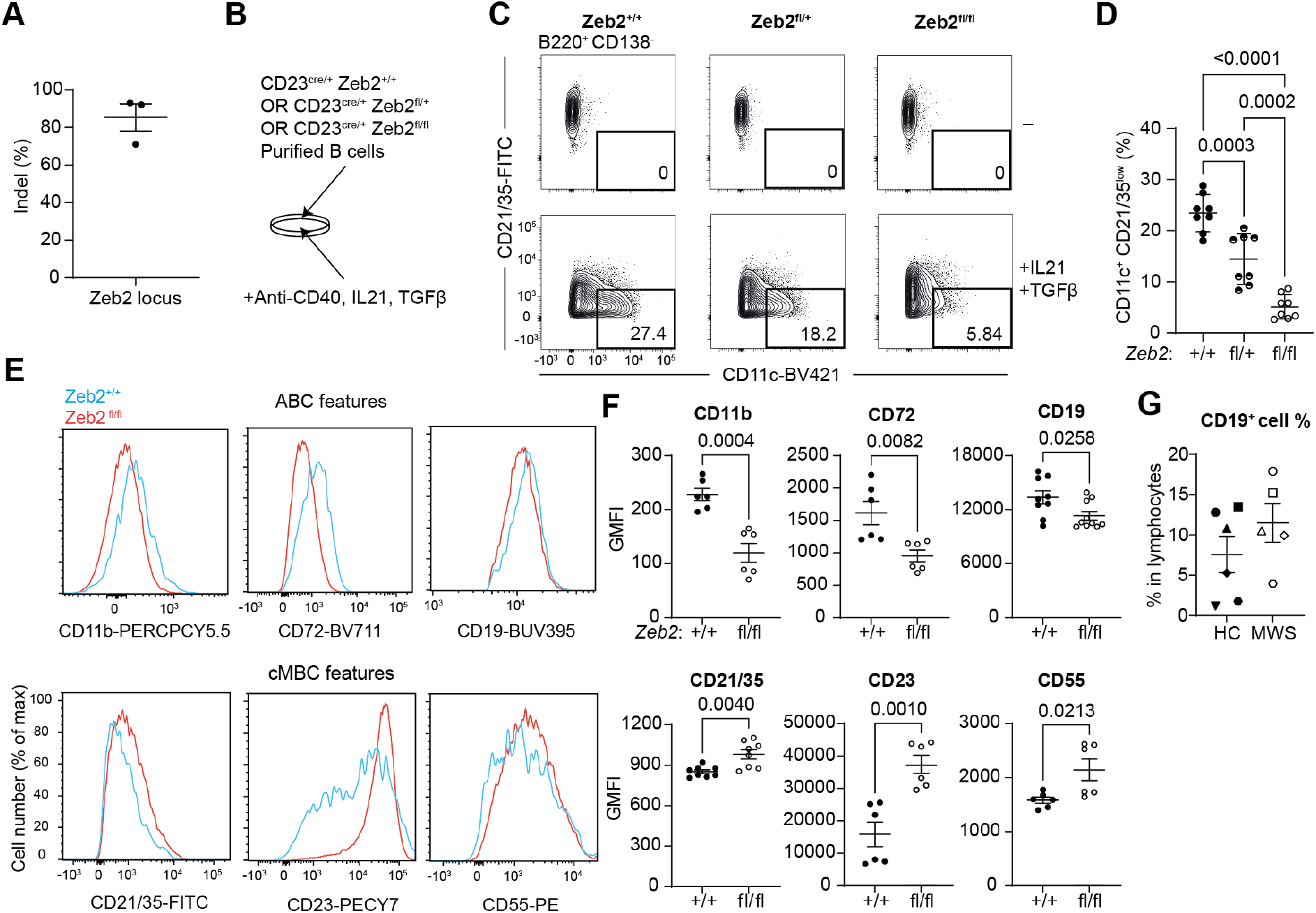
Zeb2 was required for the upregulation of ABC marker genes and suppression of cMBC marker genes. (A) Indel frequency of Zeb2 locus in Cas9 RNP transfected B cells. (B-D) MAS purified Cd23^cre/+^ Zeb2^+/+, fl/wt^ ^or^ ^fl/fl^ B cells were cultured with anti-CD40 w/wo IL-21, TGFβ for 4 days followed by FACS analysis. (B) Experiment design. (C) Representative FACS plots and (D) statistics showing the percentages of CD11c^+^ CD21/35^lo^ cells in CD138^−^ B cells. (E-F) MACS purified Cd23^cre/+^ Zeb2^+/+ or^ ^fl/fl^ B cells were cultured with anti-CD40 w/wo IL-21, TGFβ for 4 days followed by FACS analysis. (E) Representative histograms and (F) statistics showing the GMFI of ABC and cMBC markers. (G) The percentages of total B cells in lymphocytes in HC and MWS patients. The results were pooled from three independent experiments for (A), two independent experiments for (D) and two independent experiments for (F). The P values were calculated by one-way ANNOVA for (D) and student’s t test for (F).

## KEY RESOURCES TABLE

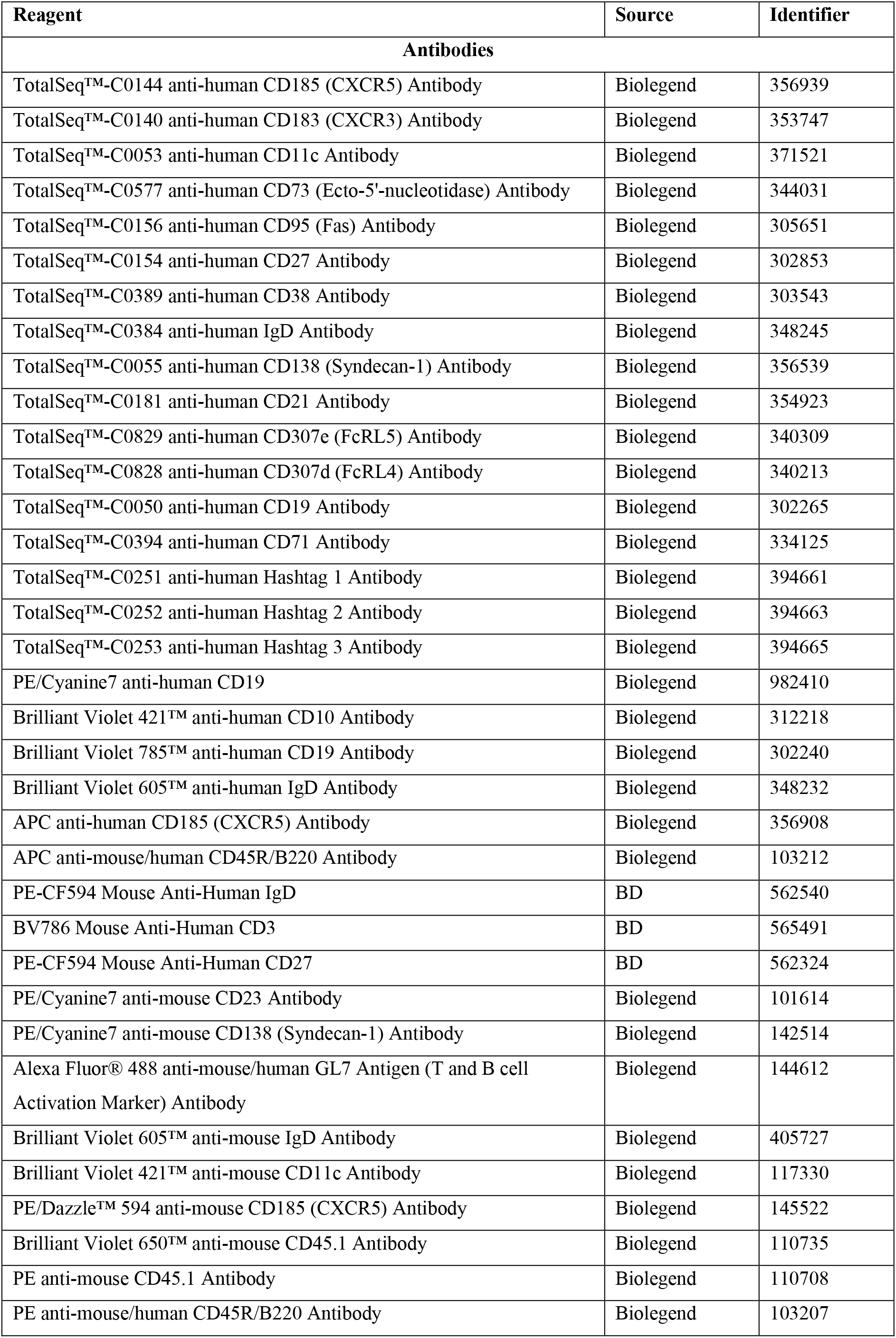
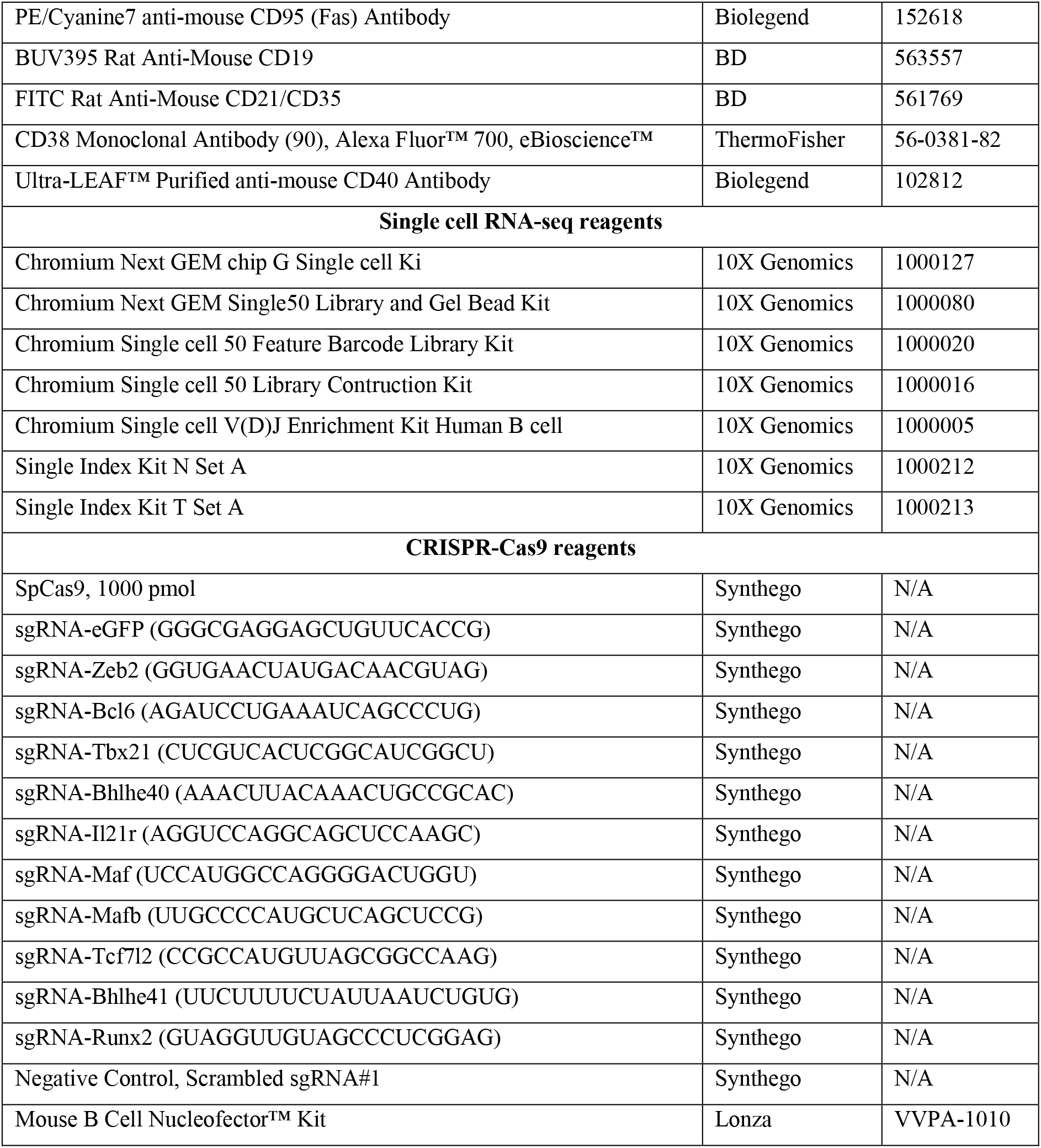

